# A 3D genome atlas of genetic variants and their pathological effects

**DOI:** 10.1101/2022.11.27.518071

**Authors:** Li Tang, Matthew C. Hill, Mingxing He, Junhao Chen, Patrick T. Ellinor, Min Li

**Author notes:** **Corresponding Author:** Min Li.

## Abstract

The spatial architecture of the genome can be categorized into distinct layers. Each layer plays a critical role in transcriptional regulation and/or genomic integrity. Alterations at any level of the 3D genome can lead to an unwanted cascade of molecular events, which may ultimately drive the manifestation of disease. However, a comprehensive atlas of the mutations and structural genetic defects that affect genome organization has yet to be compiled. Moreover, we lack a centralized resource for interpretating the pathological effects of such genetic mutations. Therefore, we curated from the literature all the pathological alterations from the chromosome level on down to single nucleotide polymorphisms (SNPs) in order to investigate these diverse genetic mutations. Using a two-phase scoring algorithm, 3DFunc, we scored the transcriptomic causality of all variants in the context of 3D genome architecture from 20 cancer and 15 normal tissues. Further, 3DFunc can identify pathological variant-gene pairs in non-oncological diseases. Finally, we constructed a web-based database, 3DGeOD (https://www.csuligroup.com/3DGeOD/home), to provide all the curated variants, genomic disruptions, as well as the scoring results derived from 3DFunc. In summary, our study constructed a 3D genome atlas of genetic variants and will serve as a valuable resource for mining the putative pathological effects of any genetic mutation.

## Background

To systematically elucidate the complex circuit of connections that exist between regulatory elements and genes, it is necessary to consider that interphase chromatin is folded in 3D in a cell-type-specific manner^1^. In recent years, chromosome conformation capture (3C) assays in combination with next-generation sequencing have provided new sights into the global organization of the genome. Since the advent of these technologies, we know that interphase genomic organization is multi-tiered. Each chromosome in the human genome occupies independent spatial territories, which play a role in the regulation of transcriptional activity and preferential positioning of loci within the nucleus^9^. The analysis of Hi-C data further refined the large-scale of territories into two sets of mega base-sized regions called “A” and “B” compartments^5^. Compartment A is enriched in the regions of high gene density, active histone markers, early replication, and open chromatin^5^. Compartment B has the opposite features compared to A, and is associated with lamina-associated domains, low transcriptional activity, late replication and, hence, are enriched in heterochromatin^5^. The topologically associated domains (TADs) are mega base pair (Mb) sized genomic regions, the TAD boundary regions contribute to the regulation of gene expression by limiting the interactions between cis-regulatory elements and target genes^10,11^. Finally, genomic loci can form specific long-range looping interactions within or across TAD boundaries, through which the regulatory elements such as enhancers and insulators play a crucial role in controlling the gene expression profile of a cell in a context-dependent manner^10,11^.

At each layer in the genome organization hierarchy, the folding patterns exhibit a complex connection to the maintenance of genomic function, and mutations that effect any of these layers may lead to disease^12,13^. Some examples include the dissolution of chromosomal territories, as observed in both breast and prostate cancers^14^. Altered genomic compartmentalization has also been reported in cancer cells. For example, by comparing normal breast cells (MCF-10A) to their cancerous counterpart (MCF-7 cells), it was found that a homogeneous switching of 12 % of all chromosomal compartments would occur^15^. Although the TAD structure is found to be a general property of the interphase chromatin across different cell types, further studies have suggested that TADs are not simple stable interactions that are formed between two permanent genomic loci, rather they are dynamic in nature^16^. The deletion of the Epha4 gene and CTCF associated boundary eliminates a TAD boundary, which causes the Epha4 promoter to interact with the Pax3 gene and drive misexpression of Pax3^17^. Genomic duplication events can affect the expression of many genes, and spurious TAD formation can also lead to human disease^18^. However, other studies have showed that interruption of TAD boundaries have no obvious impact on gene expression^19,20^.

A chromatin loop is formed when two distant genomic loci are physically closer than their intervening sequences. A classic example of long-range gene regulation involves the Shh gene, expression of which is regulated by an enhancer element ∼1 Mb away ^21^. Combining 3C techniques with genome-wide association studies (GWAS) holds great potential for identifying new putative target genes/pathways for intergenic disease-associated SNPs. Recent studies showed that most disease-associated SNPs reside within the regulatory elements and/or transcriptional factor binding sites in the non-coding regions of the genome and are likely to act through long-range chromosomal interactions. Similar effects are also shown in cases of prostate cancer^22^, breast cancer^23,24^, and in multiple other cancers^25^.

In this study, we curated the pathological alterations from the chromosome level on down to single nucleotides, including 10,789 inter-chrom translocations (ICTs), 18,863 structure variants (SVs), and 673,048 SNPs. We then analyzed the 3D disruptions caused by these curated variants, and we observed that less than 10% of ICTs interrupted territories through strong 3D interactions, and very few SVs interrupted compartments or crossed TAD structures. However, these small-scale events were found to impact gene expression significantly. Many SNPs residing within regulatory elements and/or in transcription factor binding motifs appeared to exert their effects through impacting loop strength. To investigate the transcriptional effects of variants in the context of 3D organization, we proposed 3DFunc, which combines gene expression data collected from 35 tissues and Hi-C profiles from 33 tissues to score the causality of genetic variants on transcription. Finally, we assembled a publicly available database, 3DGeOD, to provide all the curated variants, corresponding 3D layer disruptions, as well as the scoring results derived from 3DFunc.

## Results

### Disruptive Inter-chromosomal translocations (ICTs) with strong 3D interactions are highly pathogenetic

Genomic rearrangements are implicated in the pathogenesis of many types of cancer^29^. Rearrangements occur when two or more double-stranded DNA breaks are located proximally enough to fuse together (**Fig. 1a**). And recent studies have uncovered that translocation could lead to gene fusions, dysregulated gene expression, and novel molecular functions^30^ (**Fig. 1b**).

**Figure 1.**
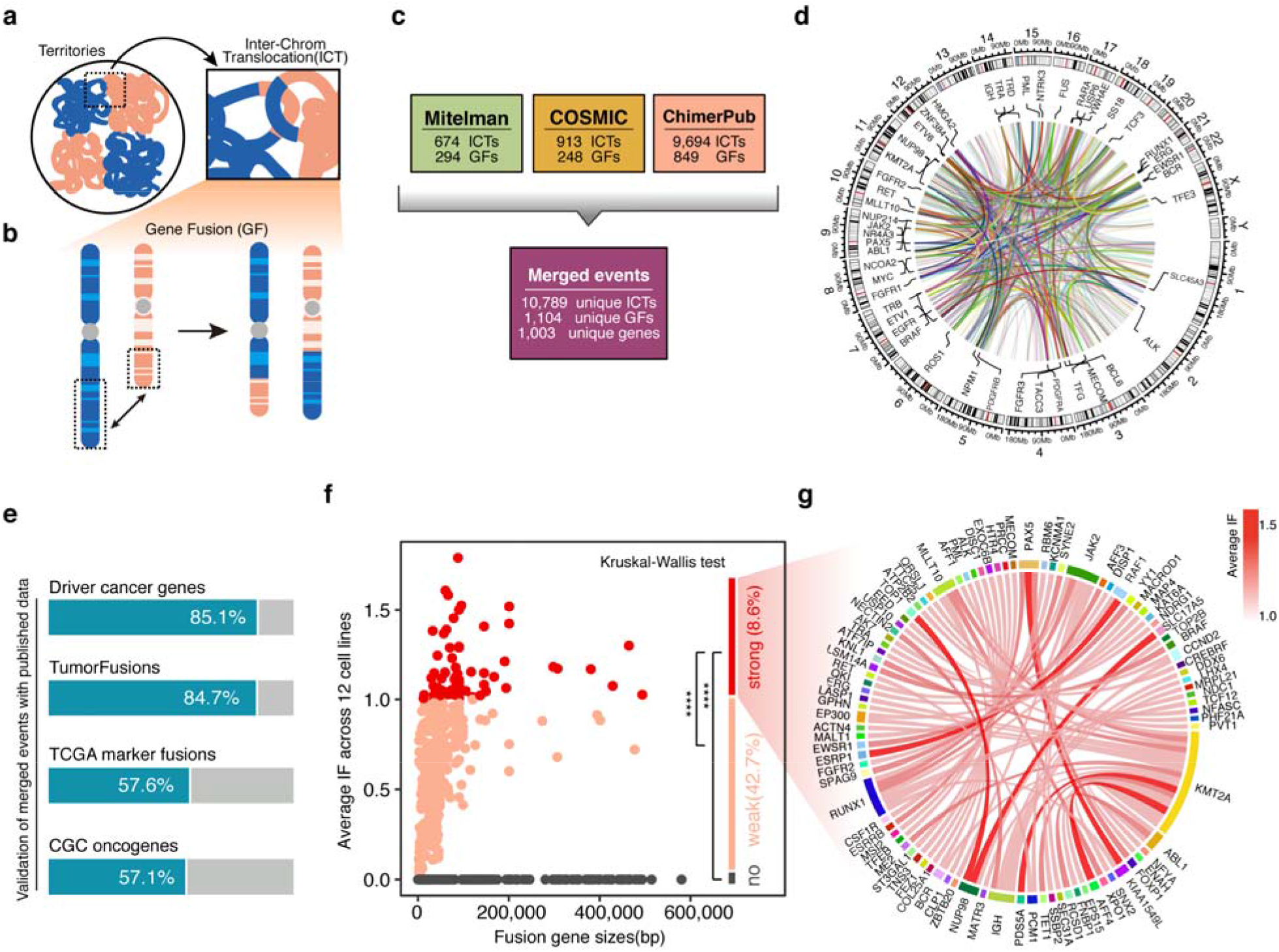
Curation of disruptive ICTs and correlation with 3D interactions. (a, b) Diagram depicting the ICTs interrupt territories, and ICTs lead to GFs. (c) The ICT events were collected from Mitelman, COSMIC, and ChimerPub. (d) The overall distribution of high frequency fusion pairs (> 10) across genome. (e) Validation of merged fusion pairs. (f) Average IF of merged fusion pairs across 12 cell lines. Difference between strong IF, weak IF and no IF was calculated with Kruskal-Wallis test with Dunn test post-hoc, **** p-value < 0.0001. (g) Circos plot of fusion pairs with strong 3D interactions.

To investigate how inter-chromosomal translocations impact gene expression through spatial proximity, we catalogued ICTs from three predominant chromosome aberration databases: Mitelman^31^, COSMIC^32^, and ChimerPub^33^ (data processing see **Methods**). Only unique events were retained, which included 10,789 unique ICTs, 1,104 unique gene fusions (GFs) and 1,003 unique genes (**Fig. 1c** and **Supplemental table 1**). There were 585 gene pairs appearing with high a frequency (> 10) across the genome (**Fig. 1d**). To validate the pathogenicity of merged events (see **Methods**), we overlaid the involved genes with driver cancer genes from DriverDBV3^34^, which showed an 85.1 % overlap. And the merged fusion events showed 84.7 % overlap with TumorFusions^35^, 57.6 % overlapping with fusions from TCGA markers, and 57.1 % overlap with oncogenes from the Cancer Gene Census^32^ (**Fig. 1e** and **Supplemental table 1**).

To correlate the pathogenic gene fusions with spatial structure of genome territories, we calculated the flexible interaction frequency (IF) between each gene pair with high-resolution Hi-C data from 12 different cell lines (calculation of flexible IF, see **Methods**). The results showed that 8.6 % of the fusion pairs had strong 3D interactions with each other, and 42.7 % had weak 3D interactions (**Fig. 1f**). Within the fusion pairs having strong 3D interactions, some of the pathogeneses have already been reported, such as *PAX5*, a transcription factor crucial for B-cell commitment and maintenance, which typically fuses with *KIAA1549L* in childhood B-cell precursor ALL (BCP-ALL)^36^. In pediatric acute leukemias, reciprocal chromosomal translocations frequently cause gene fusions involving the lysine (K)-specific methyltransferase 2A gene (*KMT2A*), specific *KMT2A* fusion partners are associated with the disease phenotype (lymphoblastic vs. myeloid), *MLLT10, PDS5A, AFF4* and so on, are common fusion partners of *KMT2A*^37^ (**Fig. 1g**). Overall, only less than 10 % of GFs interrupt chromosomal territories through strong 3D interactions, however, these GFs are highly correlated with pathogenesis.

### Structural variations (SVs) don’t frequently interrupt compartment switching but do cause significant changes in gene expression

Next-generation sequencing (NGS) has revealed the enormous structural complexity of the human genome^38,39^. Structural chromosomal rearrangements referred as structural variation (SV), contribute to a large extent of the genetic diversity in the human genome and thus are of high relevance for cancer genetics, rare diseases, and evolutionary genetics^40^. Recent studies have shown that SVs can alter the copy number of regulatory elements or modify the 3D genome by disrupting higher-order chromatin organization such as compartments and TADs ^41,42^. As a result of these position effects, SVs can influence the expression of genes distant from the SV breakpoints, thereby causing disease. The impact of SVs on the 3D genome and on gene expression regulation need to be considered when interpreting the pathogenic potential of such genetic variations^43^.

We examined 18,863 pathogenic SVs from Clinvar^44^, which included copy number gain, copy number loss, deletion, duplication, and insertion. We then filtered these SVs for length, selecting those with a size of 10 kb to 10 Mb for the subsequent analysis (**Supplemental Fig. S1A**). The filtered SVs were mapped to A/B compartments, four types of compartment interruptions including: A-A (both breakpoints within A compartments), B-B (both breakpoints within B compartments), A-B (left breakpoint within A compartment and right breakpoint within B compartment), B-A (left breakpoint within B compartment and right breakpoint within A compartment), in which A-A and B-B were defined as stable compartment interruptions, A-B and B-A were defined as switching compartment interruptions (for detection of compartments, see **Methods**). We found that two breakpoints of SV occurred more frequently within regions having no compartmental changes, especially in the type of A-A. While the SVs that did cause compartment switching occurred in less than 20% of all SVs, such as A-B or B-A (**Fig. 2a-2b** and **Supplemental table 2**). Next, we calculated the flexible interaction frequency (IF) (see **Methods**) for all four possible types of compartmental disruptions, and the stable compartment interruptions showed more 3D interactions than switching types (**Fig. 2c** and **Supplemental table 2**). In addition, we observed that the SVs mapped to switching compartments involved more dosage sensitivity genes, consistent with a previous study which found that altered compartmental states could lead to transcriptional activity change^45^ **(Fig. 2d** and **Supplemental table 2)**. Overall, most of the SVs we interrogated interrupted stable compartments, which also possessed high levels of 3D interactions. Although only a small portion of SVs caused compartment switching, these SVs involved greater transcriptional activity changes through less 3D interactions.

**Figure 2.**
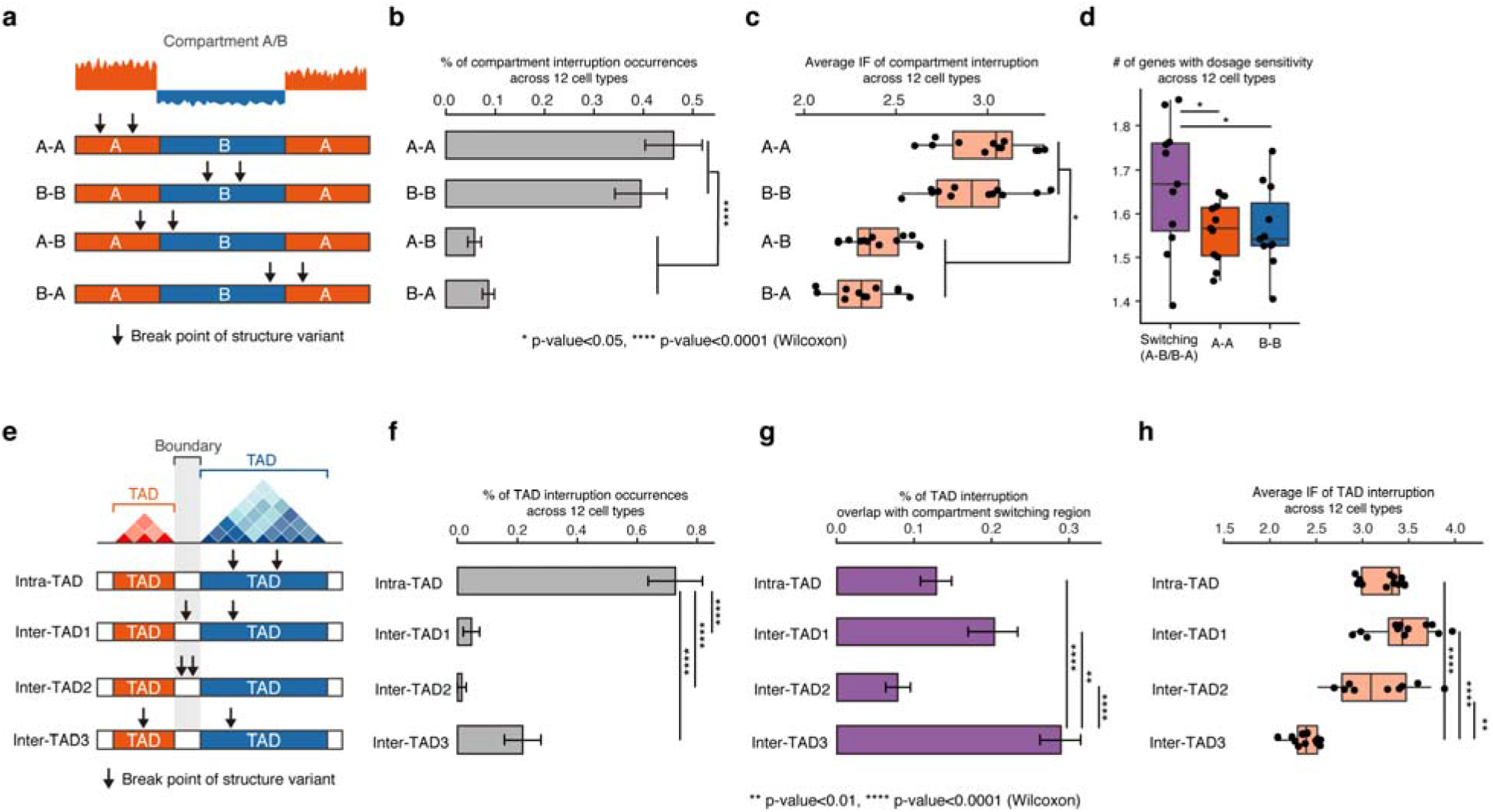
Interruptions of compartments and TADs caused by SVs. (a) Four types of compartment interruptions: A-A (both breakpoints within A compartments), B-B (both breakpoints within B compartments), A-B (left breakpoint within A compartment and right breakpoint within B compartment), B-A (left breakpoint within B compartment and right breakpoint within A compartment). (b) Percentage of compartment interruption occurrences across 12 cell types. (c) Average IF of compartment interruption across 12 cell types. (d) Number of genes with dosage sensitivity across 12 cell types for the interruptions in switching and stable compartment. (e) Four types of SVs interrupt TADs: intra-TAD (both breakpoints located within the same TAD), inter-TAD1 (with one breakpoint locate in boundary and the other in TAD), inter-TAD2 (both breakpoints located within the boundary), and inter-TAD3 (two breakpoint locate in different TADs). (f) Percentage of TAD interruption occurrences across 12 cell types. (g) Percentage of TAD interruptions overlap with compartment switching regions. (h) Average IF of TAD interruption across 12 cell types. P-value were calculated by Wilcoxon test, * p-value < 0.05, ** p-value < 0.01, *** p-value < 0.001, **** p-value < 0.0001.

Recent studies have reported that structural variation can induce dramatic changes in TAD organization^17,18^. We mapped SVs to TAD boundaries (for detection of TADs, see **Methods**), and categorized the predicted interruptions into four categories: intra-TAD (both breakpoints located within the same TAD), inter-TAD1 (with one breakpoint locate in boundary and the other in TAD), inter-TAD2 (both breakpoints located within the boundary), and inter-TAD3 (two breakpoints locate in different TADs) (**Fig. 2e**). Among the four types of interruptions, intra-TAD occupied the largest proportion, followed by inter-TAD3 (**Fig. 2f** and **Supplemental table 2**). We then mapped four types of TAD interruptions to the switching compartments regions. Inter-TAD3 had the highest overlapping percentage with switching compartments, indicating that the inter-TAD3 interruptions potentially correlated with greater transcriptional activity changes (**Fig. 2g**). We then calculated the flexible IF for SVs in all four types of TAD interruptions. Inter-TAD3 showed the lowest 3D connection frequency (**Fig. 2h** and **Supplemental table 2**). Overall, fewer SVs occur in inter-TAD than intra-TAD, and the type of inter-TAD3 correlates more with compartment switching, but also displays lower amounts of 3D genomic interactions.

### Analysis of disease-associated single nucleotide polymorphisms (SNPs)

Many human genes are regulated by at least one regulatory element that influences the transcriptional state of their respective promoters. High-resolution 3C data strongly suggests the presence of regulatory chromosomal loops as a means of communication among the local and/or distal regulatory elements with promoters to control gene expression ^10^. In recent studies, the presence of a SNP either at the enhancer side or at the promoter element of an enhancer-promoter interaction could cause altered expression of the corresponding gene ^21,46,47^. Expression quantitative trait loci (eQTL) mapping has paralleled the adoption of GWAS for the analysis of complex traits and disease in humans. Under the hypothesis that many GWAS associations tag non-coding SNPs with small effects, and these SNPs exert phenotypic control by modifying gene expression, it has become common to interpret GWAS associations using eQTL data 48,49.

In this study, we extracted the fine-mapped SNP-gene loops from recomputed datasets of the eQTL Catalogue ^50^. We considered 9 tissues in the eQTL Catalogue which had corresponding high-resolution Hi-C data available (for processing of fine-mapping eQTL data, see **Methods**). The number of credible sets (CSs) ranged from 1,034 to 7,269, and the number of SNPs ranged from 31,776 to 282,005 per tissue (**Fig. 3a** and **Supplemental table 3**). The size of CSs between different tissues showed a similar distribution (**Fig. 3b** and **Supplemental table 3**). We observed that the CSs with a size larger than 2,000 were sparse and had a higher average r^2^ value across all the tissues (**Supplemental Fig.S2**). The posterior inclusion probability (PIP) was computed by SuSiE ^51^ and was utilized to calculate the causality of variants. Overall, there were 3,596 SNPs with a high PIP ≥0.95 across all the tissues (**Fig. 3c** and **Supplemental table 3**), and blood observed the most SNPs with high PIP (≥0.95) within these tissues (**Supplemental Fig.S3-S4** and **Supplemental table 3**). To interpret the pathological effect of SNPs in the context of 3D genome architecture, we computed the flexible IF for each SNP-gene loop. The results showed that SNP-gene loops with a higher PIP had more 3D interactions (**Fig. 3d** and **Supplemental table 3**). Recent studies have uncovered that many disease-associated SNPs reside within regulatory elements and/or are enriched in transcriptional factor binding motifs in the non-coding region of the genome and exert their effects through long-range chromosomal interactions ^52,53^. Thus, here we examined whether SNPs caused non-coding region changes or affected RefSeq and UCSC genes, then annotated them with ENCODE DNase I hypersensitive sites (DHSs). And we employed SNP2TFBS ^54^ to investigate the transcriptional factor binding sites (TFBS) associated with each SNP (see **Methods**). We found 144,052 sites with TFBS matches, and the number of affected TFBS ranged from 1 to 11 within each site (**Supplemental Fig.S5**). Genomic region annotation showed that most of the SNPs were located within non-coding regions (89.2 %), and 30.5 % of which resided in DHS regions and/or within TFBS (**Fig. 3e**). SNPs in coding regions had a higher percentage of PIP ≥0.95 than non-coding regions, and the SNPs in DHS peaks and/or within TFBS were observed to have a higher PIP than other regions (**Fig. 3f** and **Supplemental table 3**). The IF calculation results of the different SNP types showed that there were more 3D connections between SNP-gene pairs when SNPs were located within DHS regions and/or within a TFBS in both coding and non-coding regions (**Fig. 3g** and **Supplemental table 3**).

**Figure 3.**
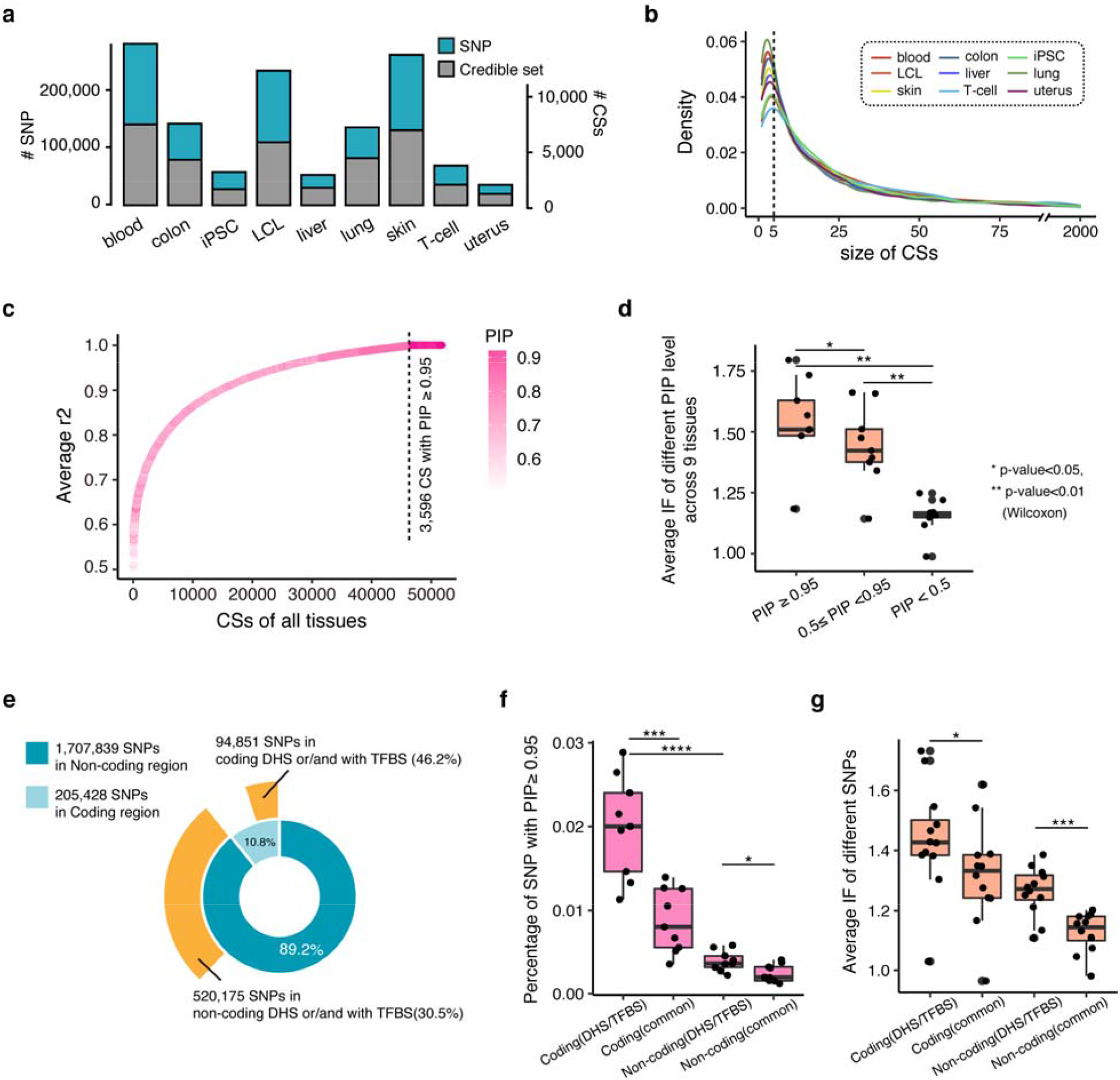
The causality and localization preference of SNP-gene loops. (a) The number of SNPs and CSs. (b) The size distribution of CSs. (c) PIP of all SNPs ranked by average r2. (d) Average IF of different PIP levels across 9 tissues. (e) Percentage of SNPs in coding, non-coding, DHS region or/and with TFBS. (f) Percentage of SNPs with PIP≥0.95 in different types of regions. (g) Average IF of SNPs in different types of regions. P-value were calculated by Wilcoxon test, * p-value < 0.05, ** p-value < 0.01, *** p-value < 0.001, **** p-value < 0.0001.

Overall, SNP-gene loops with a higher PIP had more 3D interactions, and disease-associated SNP-gene loops localized preferentially within DHSs containing elements or within a predicted TFBS.

### 3DFunc: A two-phase scoring algorithm for detecting the variants that affect gene function through long-range genomic interactions

To enlarge the testing datasets, we identified ICTs, SVs, and SNPs from the Pan-cancer analysis of whole genomes (PCAWG) ^27^ (**Supplemental Fig.S6**., **for** Calling variants see **Methods**). Firstly, we collected gene expression data from PCAWG, and high-resolution Hi-C data from 4DN ^26^, then ranked the predicted variants with downstream expression changes and IF changes individually (for processing of gene expression and Hi-C data, see **Methods**). The verified causal variants didn’t show a preferable top ranking with a single data type, indicating the causality of variants was complex and cannot be characterized with only a single data type (**Fig. 4a** and **Supplemental table 4**). To quantify the effect of variants in the context of 3D interactions, we built a two-phase scoring algorithm, 3DFunc, which integrates both high-resolution Hi-C profiles and gene expression data to fit a nonlinear least square curve (**Fig. 4b**, see **Methods**). We then used 3DFunc to score the predicted variants and calculate the percentage of verified causal variants within the top 10 % of our 3DFunc ranked variants, which showed that 3DFunc ranked verified causal variants preferably (**Fig. 4c-4d** and **Supplemental table 4)**.

**Figure 4.**
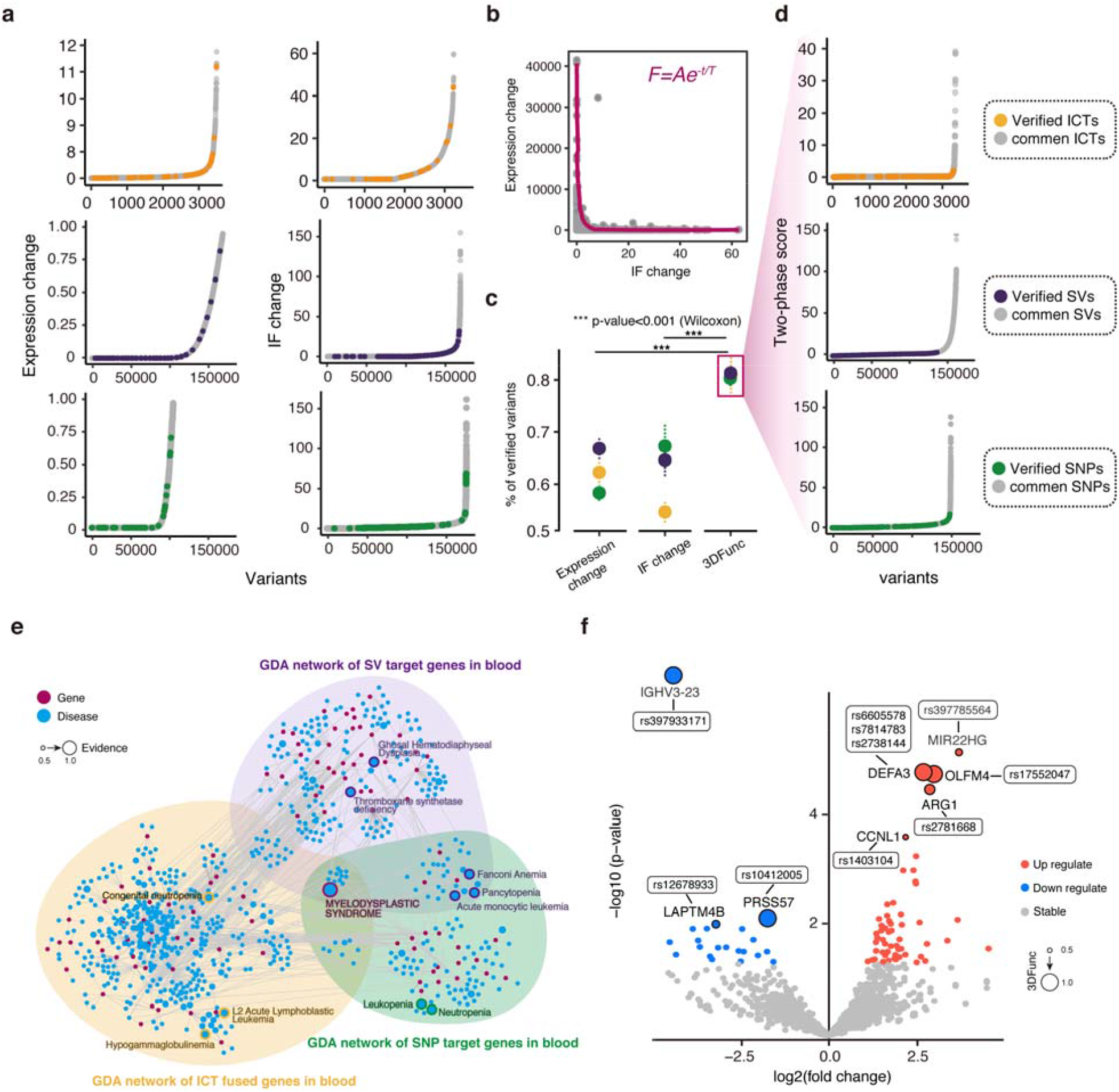
3DFunc predicts the transcriptional effects of diverse sets of genetic variants. (a) Rank variants with expression change or IF change individually. (b) 3DFunc integrates both high-resolution Hi-C profile and gene expression data to fit the nonlinear least square curve. (c) The percentage of verified causal variants within the top 10% ranking parts, p-value was calculated by Wilcoxon test, *** p-value < 0.001. (d) Rank variants with 3DFunc scores. (e) GAD network of ICTs, SVs, and SNPs in blood tissue. (f) Differential expression analysis for TCGA MDS data with 3DFunc scoring, p-value < 0.5 and the absolute value of log2(fold change) larger than or equal to 1 were used as the significant threshold.

Next, we extracted the ICT-gene, SV-gene, and SNP-gene pairs from 5 common tissues and calculated the significance of 3DFunc scoring (chi-square test). We then overlayed the driver mutations from PCAWG with the variants in these common tissues. The variants in blood showed both the most significant 3DFunc scores (p-value < 0.05) and the highest driver mutation rate. (**Supplemental Fig.S7**). To investigate the pathogenesis of 3DFunc scores, we analyzed the Gene Disease Association (GDA) network for the variants target genes (for construction of the GDA network, see **Methods**). Myelodysplastic Syndrome (MDS) was observed in three types of variants with the highest evidence of disease risk (**Fig. 4e, Supplemental Fig.S8** and **Supplemental table 5**). We then used gene expression data of both control and MDS samples from TCGA to filter the variants target gene(s), which showed 85 genes were significantly differentially expressed. And some of them were observed with high 3DFunc scores, such as *IGHV3-23*, which is one of the most frequently mutated IGHV genes in the chronic lymphocytic leukemia series ^55^. Further, *DEFA3*, which is highly expressed in neutrophils, was observed interacting with three pathological SNPs (rs6605578, rs7814783, and rs2738144) with high 3DFunc scores (**Fig. 4f** and **Supplemental table 6**. For analysis of TCGA MDS data, see **Methods**).

To further verify the effectiveness of 3DFunc, we extracted the predicted SVs and corresponding interactions within the *WNT6*/*EPHA4/PAX3* locus. We analyzed H1 cell Hi-C profiles^56^ to annotate the TAD structures at this locus. The 3DFunc score of W-SV-3 (the third nearest SV of WNT6) was the highest within all the WNT6 related SVs, which was close to the TAD boundary. The region near the *WNT6* TAD boundary when duplicated will misplace the *WNT6* gene closer to an enhancer element causing its misexpression^17^. For the SVs of *EPHA4*, three 3DFunc scores were larger than 0.5, indicating the causal effect of the upstream enhancer region of *EPHA4*. For comparison, we also showed the scoring of the Caviar Probability, a measure of causal variants in associated regions^57^, which didn’t display higher scores for the SV-*WNT6* and SV-*EPHA4* pairs (**Supplemental Fig.S9**).

### Application of 3DFunc to four common diseases

For various common diseases, interpretating the molecular mechanism of any given noncoding variant remains a challenge. And given the functionality of our tool and scoring procedure, we sought to apply our algorithm to a diverse set of disease classes to further demonstrate its utility for predicting the effects of genetic mutations. Here we studied four common diseases with major public health impact: cardiovascular disease, type 2 diabetes (T2D), inflammatory bowel disease (IBD), and breast cancer. RNA-seq expression and Hi-C data derived from the heart, pancreatic islet, intestine, and breast tissues were used as input to calculate the 3DFunc scores individually (see **Methods**).

To investigate the pathogenesis of detected variant-gene pairs, we calculated gene pathway enrichment with DisGeNET ^58^, which showed related disease enriched categories in the respective tissues (**Fig. 5a** and **Supplemental table 7**. For gene list enrichment, see **Methods**). We then used genome-wide polygenic score (GPS) to evaluate the risk of variants with clinical significance. In a previous study, several candidate GPSs were calculated based on summary statistics and imputation from large GWAS datasets with participants of primarily European ancestry ^59^. Here we mapped the GPS to our calculated 3DFunc scores by splitting the variants into independent percentile groups for each disease (see **Methods**), and the average GPS of each group was found to increase according to the disease-specific 3dToFuc percentile, which highlighted the clinical significance of our top-ranking variants (**Fig. 5b** and **Supplemental table 8**). For the four chosen common diseases, the SNP-gene pairs with the top 1 % 3DFunc scoring were selected, in which 1193 for cardiovascular disease, 198 for T2D, 403 for IBD, and 10 for breast cancer, respectively. And we checked the pairing counts of SNP and gene, which showed an average 1.31 SNPs paired to one gene and averaged 1.23 genes paired to each SNP across the four diseases we inspected (**Fig. 5b**).

**Figure 5.**
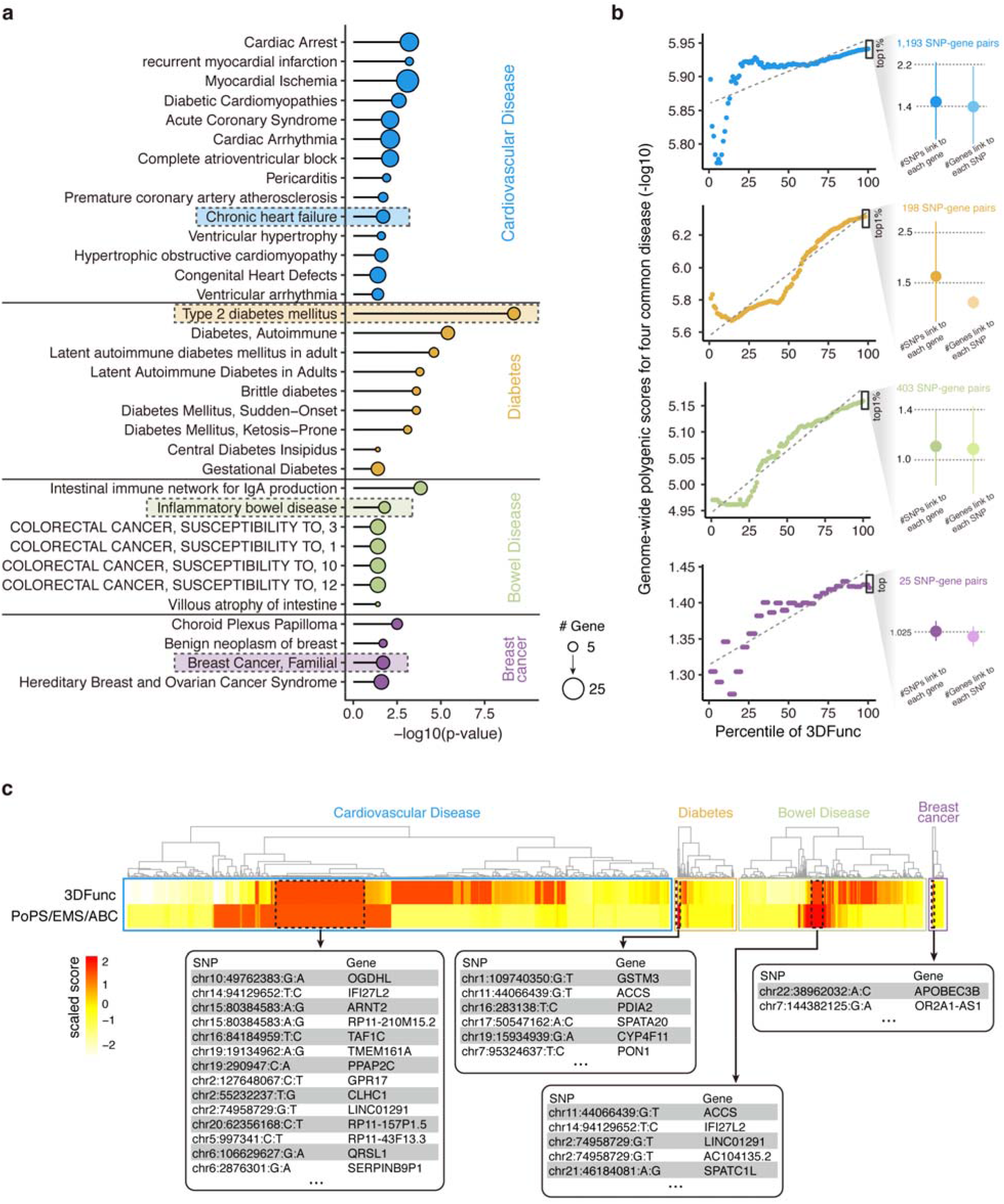
Application of 3DFunc to four common diseases. (a) GO enrichment of gene list with DisGeNET in heart, pancreatic islet, intestine, and breast tissues. The size of dot represented the number of genes enriched in each disease. (b) Average GPS according to the disease-specific 3dToFuc percentile, the pairing counts of SNP and genes in four common diseases. The regression line was fitted with a linear model. (c) Mapping of 3DFunc scores to PoPS/EMS/ABC, and the consistent highly scored pairs were showed.

As some fine-mapping studies have been proposed to identify the putative causal eQTL, here we curated the fine-mapping results of expression modifier score (EMS) in 49 GTEx tissues^60^, and the Polygenic Priority (PoPS) Score in 18 diseases with publicly available GWAS summary statistics and 95 complex traits from the UK Biobank^61^, as well as the activity-by-contact (ABC) scores in 72 diseases and complex traits^62^. We mapped our top-ranking variant-gene pairs with the fine-mapping results, and extracted the consistent highly scored pairs (**Fig. 5c** and **Supplemental table 8**). From this we identified 122 consistent pairs for heart disease, 8 for T2D, 35 for IBD, and 3 for breast cancer, the validation of these putative causal eQTLs highlights the ability of 3DFunc to detect causal variant-gene pairs.

### A 3D genome atlas of genetic variants and their pathological effects

In the above work, thousands of pathological genetic mutations were curated, and we analyzed the topological genomic disruptions caused by these curated variants (**Figure 6a**), in which 1104 ICTs interrupted territories, 5002 SVs correlated with compartments switching events, 7654 SVs disrupted TAD boundaries, and 3033 SNPs disrupted loops. To interpret how these variants impact gene expression through 3D interactions in a quantitative manner, we generated a two-phase scoring algorithm known as 3DFunc, which combines 33 Hi-C datasets from the 4DN data portal^26^, gene expression data of 20 cancer tissues derived from the Pan-cancer analysis of whole genomes (PCAWG)^27^ and 15 normal tissues taken from the GTEx portal^28^. 3DFunc employs a nonlinear least square curve to measure the effect of genetic variants on long-range gene regulation and genomic architecture. To demonstrate the effectiveness of 3DFunc, more ICTs, SVs and SNPs were identified from the PCAWG datasets, and scored with 3DFunc. Finally, we integrated all of the curated data and the calculated 3DFunc scoring results to the 3DGeOD database, in order to provide a atlas of genetic variants and their pathological effects for further research, which is available at https://www.csuligroup.com/3DGeOD/home. (**Figure 6b**).

**Figure 6.**
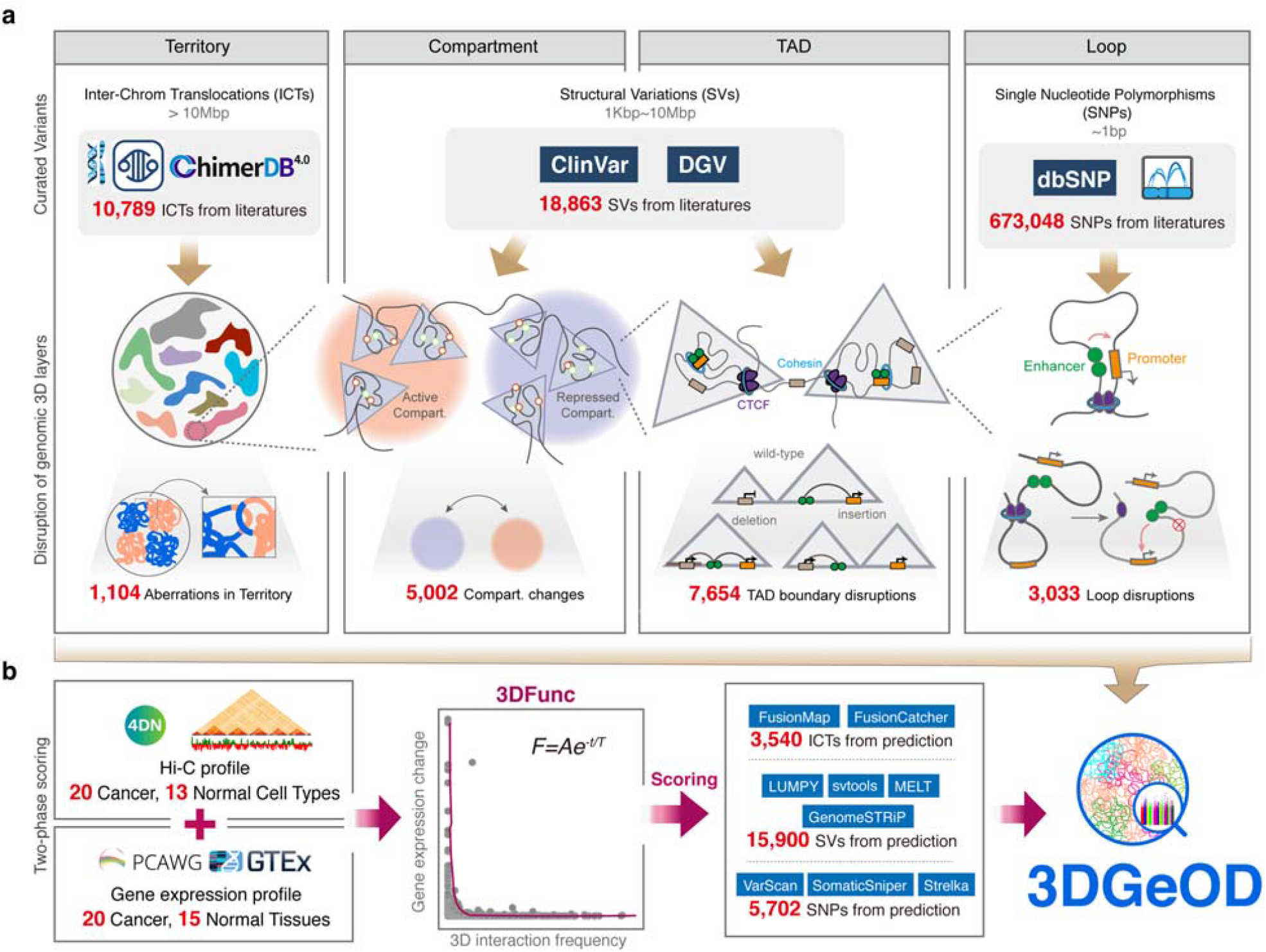
Design of 3DGeOD. (a) 10789 ICTs, 18863 SVs and 673048 SNPs were curated from literatures, in which 1104, 5002, 7654, and 3033 events were detected disrupting the corresponding 3D architecture layers. (b) Hi-C profile from 33 tissues and gene expression data from 35 tissues were combined with a nonlinear least square curve to generate two-phase scoring algorithm 3DFunc. The predicted variants were scored with 3DFunc. The database 3DGeOD was constructed with the literature curated data and the 3DFunc predictions.

## Discussion

In recent years, chromosome conformation capture (3C) assays in combination with high-throughput sequencing, such as Hi-C, has provided new sights into the global organization of the genome. Since the advent of these techniques, we know that interphase chromosomes are folded into four layers of hierarchical structures, including territory, compartment, TAD, and loop. The alterations in any of these layers can lead to disease or phenotype, thus we curated the variants and mapped them to the 3D layers to investigate the pathological effects.

For territory, ICT could lead to gene fusions and dysregulated gene expression through spatial proximity, we observe that the fused gene pairs with more 3D interactions tend to corelate with pathogenesis. It was reasonable, as DNA FISH studies have uncovered that the frequency of translocations between two chromosomes is related to their spatial proximity in interphase nuclei ^63,64^. Consistent with this finding, the 3D interaction counts from Hi-C profile between whole chromosomes were found to exhibit a strong correlation with the frequency of translocation, suggesting that spatial genomic proximity precedes translocation ^65^.

For compartment, SV influence the expression of genes distant from the SV breakpoints and caused disease. In this study, most of the pathogenic SVs interrupt stable compartments, which also involve more 3D spatial connections. Although small part of SVs interrupt switching compartments, these SVs impact more gene expression through less 3D interactions. For TAD, fewer SVs occur inter-TAD, and the type of inter-TAD3 (two breakpoint locate in different TADs) correlated more with switching compartment and had less 3D interactions. Previous studies have demonstrated that compartments and TADs were highly dynamic in nature and changes occurred in accordance with lineage and cell-type specificity^66^. What’s more, the altered compartmentalization and interruptions of TAD boundaries have been reported in disease and complex traits ^15,18^. Which explained why we observed that the SVs occurred in switching compartment and inter-TAD structures impacted more dosage gene expression. In addition, less 3D interactions were detected near these regions, which was possibly caused by its dynamic characteristics.

For loop, we observe that the SNP-gene loops locating in the DHS regions or with TFBS are observed higher 3D interaction frequencies and with higher causality. Consistent with the previous cancer studies that most of the disease associated SNPs reside within the regulatory elements and/or enriched in transcriptional factor binding motifs in the non-coding region of the genome and exerts effects through long range chromosomal interactions^22,23^.

Therefore, based on the 3D genome atlas of variants we depicted, we employed the nonlinear least square curve to fit the gene expression and 3D contacts, and proposed a two-phase scoring algorithm, 3DFunc, to identify the variants which employ 3D interactions to affect function in pathogenesis. 3DFunc takes only RNA-seq and Hi-C data as input, the causality of variant-gene pairs will be scored and output. We use 3DFunc to 20 cancer and 15 normal tissues to obtain 3540 ICT, 15900 SV-gene, and 533349 SNP-gene pairs with pathological scores. The application of 3DFunc to four common diseases reveals its effectiveness of detecting causal variant-gene pairs.

## Conclusion

3C assays in combination with high-throughput sequencing have provided new sights into the global organization of the genome, and the genomic mutations of different scales may lead to disease or complex traits. Here we curated pathological ICTs, SVs and SNPs from literatures to investigate the corresponding 3D layers interruptions, as well as understand their pathological effects. Then a two-phase scoring algorithm, 3DFunc, was proposed to detect the causal variant-gene pairs in the 3D context, and we applied 3DFunc to four common diseases to demonstrate its effectiveness. Finally, we constructed a database, 3DGeOD, which included thousands of curated genetic mutation data along with the corresponding 3D layer interruptions, and the scoring results of 3DFunc.

## Methods

### Curation of ICTs

The ICTs events were collected from three predominant chromosome aberration database: Mitelman^31^, COSMIC^32^, and ChimerPub^33^. For the data from Mitelman database (https://mitelmandatabase.isb-cgc.org/), only the structural aberrations of inter-chromosome were retained, and the karyotype of which were converted to chromosome coordinates with CytoConverter^72^. For the data from COSMIC (https://cancer.sanger.ac.uk/cosmic), the structural genomic rearrangements were downloaded, and the inter-chromosomal translocations were filtered out. For the data from ChimerPub (https://www.kobic.re.kr/chimerdb/chimerpub), the translocation data with known chromosome positions were retained. Then the all the ICTs from these three databases were merged and removed duplications, 10789 ICTs were retained in total. 1104 gene fusion events were extracted from the filtered unique ICTs. All the chromosome coordinates were extracted under the reference of hg38.

### Curation of SVs

The pathological SVs were collected from Clinvar ^44^ (release of 2021-08-13) under the reference of hg38 (https://ftp.ncbi.nlm.nih.gov/pub/dbVar/sandbox/dbvarhub/hg38/), we filtered the SVs with the percentage of larger than 1%, the filtered SVs included copy number gain, copy number loss, deletion, duplication, and so on. For the subsequent analysis, 18863 SVs with length of 10k to 10m bp were retained.

### Curation of SNPs

The SNPs were collected from dbSNP ^73^ (release of 2021-05-25) under the reference of hg38 (https://www.ncbi.nlm.nih.gov/snp/). To narrow the scope to pathological SNPs, we downloaded the fine mapping eQTL data from the recomputed datasets of eQTL Catalogue ^50^ (https://www.ebi.ac.uk/eqtl/Data_access/), the credible sets of the same tissue from different studies were merged and the duplicated SNP-gene pairs were removed. Then we overlayed the SNPs in dbSNP with the fine-mapping SNPs, only SNPs with the highest posterior inclusion probability (PIP) within the credible set were retained for the subsequent analysis.

### Disruption of 3D layers

For the layer of territory, the filtered unique gene fusions from curated ICTs were regarded as the disruptions of territory, and the cytoband of two fused genes were regarded as the locations where aberrations occurring.

For the layer of compartment, we firstly used high-resolution Hi-C datasets (>1 billion reads) of 12 cell types from 4DN data portal^26^ to assign active (A) or inactive (B) compartments genome-wide by FAN-C^74^ with the resolution of 1Mb, and the genome sequence of hg38 was applied to calculate the average GC content, as GC content has previously been shown to correlate well with compartmentalization^74^. Then we overlayed the curated SVs with the compartments of 12 cell types individually, the spanning locations of SVs were regarded as the disruptions of compartments. There were four different disruption types, A-A (both breakpoints within A compartments), B-B (both breakpoints within B compartments), A-B (left breakpoint within A compartment and right breakpoint within B compartment), B-A (left breakpoint within B compartment and right breakpoint within A compartment).

For the layer of TAD, we used high-resolution Hi-C datasets (>1 billion reads) of 12 cell types from 4DN data portal to call the insulation domains by FAN-C with the window size of 1Mb. Since the downloaded Hi-C file has been normalized with ICE^75^, here we didn’t perform any normalization. We overlayed the curated SVs with the TADs of 12 cell types individually, the spanning locations of SVs were regarded as the disruptions of TADs. There were four interruption types: intra-TAD (both breakpoints located within the same TAD), inter-TAD1 (with one breakpoint locate in boundary and the other in TAD), inter-TAD2 (both breakpoints located within the boundary), and inter-TAD3 (two breakpoint locate in different TADs)

For the layer of loop, the curated SNPs were assigned to the target genes with fine-mapping eQTL data, the chromosome regions near SNPs were extended to 1kb on both sides, then the extended locus-gene pairs were regarded as loops, and the SNPs were regarded as the interruption within loops.

### Calculation of flexible IF

The flexible IF used Hi-C matrix to measure the interaction frequency between two specific genomic loci. To make the calculation fit different scale of genomic length, we used a flexible strategy to determine the window size of interaction frequency. We defined two candidate regions as *C*_*a*_ and *C*_*b*_, respectively. The start point of *C*_*a*_ was *S*_*a*_, the end point of *C*_*a*_ was *E*_*a*_, the start point of *C*_*b*_ was *S*_*b*_, the end point of *C*_*b*_ was *E*_*b*_. The Hi-C files created by juicer ^75^ contained several pre-built matrices with different resolutions, such as 1000bp, 5000bp, 10000bp, 50000bp, 100000bp, and 500000bp. For the candidate regions *C*_*a*_ and *C*_*b*_, we can determine the nearest resolution *R*_*a*_ and *R*_*b*_, respectively. The window size for calculating interaction frequency was set to *W*_*ab*_ = min {*R*_*a*,_ *R*_*b*_}. The interaction frequency between *C*_*a*_ and *C*_*b*_ was calculated by Straw^75^, which was represented by *IF* (*S*_*a*_:*E*_*a*_,*S*_*b*_,:*E*_*b*_, *W*_*a,b*_}.

Since most of the Hi-C files were low-resolution, which made the IF sparse, here we extended the candidate regions as background to calculate the flexible IF. The extended start point of *C*_*a*_ was 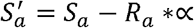, the extended end point of *C*_*a*_ was 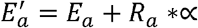, the extended start point of *C*_*b*_ was 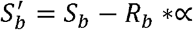, the extended end point of *C*_*b*_ was 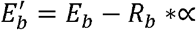. The extension coefficient was ∝ set to 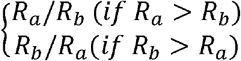. Then the flexible IF was calculated by

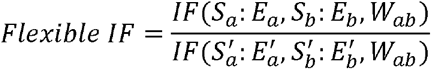

For the ICTs events, the locus of two fused genes were extracted as candidate regions for calculation. For the SVs events, all the SVs-gene pairs within the same TAD structure were considered as the candidate regions. For the SNPs-gene pairs, the chromosome regions near SNPs were extended to 1kb on both sides, then the extended locus-gene pairs were regarded as candidate regions.

### Validation of merged ICTs

To validate the pathogeneses of merged ICTs, we collected the driver cancer genes from DriverDBV3^34^ (http://driverdb.tms.cmu.edu.tw/download), then overlayed the genes involving in the ICTs with the driver cancer genes, and calculated the overlapping percentage. For the gene fusions from the whitelist of TumorFusions^35^ (https://tumorfusions.org/), the fusions from TCGA marker papers, and the TCGA fusions dataset^76^, we overlayed the curated gene fusions with them regardless the fusion directions, then calculated the overlapping percentage.

### Genomic annotation of SNPs

To investigate the 3D interaction characteristics of SNPs within different genomic regions. We annotated the curated pathological SNPs to coding regions with the coordinates of known genes from UCSC^77^ (release of 2021-11-16) and RefGene^78^ (release of 2021-03-01), and the noncoding SNPs did not locate in a coding sequence or within 10 bp of a splice site annotated in the RefGene. And we annotated the SNPs with high resolution maps of DHSs data^79^. Then we employed SNP2TFBS ^54^ (https://ccg.epfl.ch/snp2tfbs/) to investigate the transcriptional factor binding sites (TFBS) associated with coding and noncoding SNPs respectively. The enrichment of TF and the number of affected TFBS were calculated by SNP2TFBS.

### Process of PCAWG and GTEx data

For PCAWG gene expression data, a total of 1,359 samples in 20 cancer tissues were used, the reads were aligned with the alignment tools of TopHat2^80^ (release of v2.1.1) and STAR^81^ (release of v2.7.10a). Read counts to genes were calculated using Htseq-count^82^ with GFT file from GENCODE human release v38. Then counts were normalized using FPKM normalization and upper quartile normalization (FPKM-UQ). The final expression values are given as an average of the TopHat2 and STAR-based alignments.

For GTEx data, the expression of 15 healthy tissues in 3,274 samples from GTEx (phs000424.v4.p1) were analyzed with the same pipeline as PCAWG gene expression data to calculate the FPKM value for genes. The calculated expression data of each sample was assigned a unique aliquot.

### Identification of ICTs

The gene fusions of ICTs were identified from the alignment files of PCAWG with FusionMap^83^ (release of 2015-03-31) and FusionCatcher^84^ (release of v1.33). FusionMap was employed to detect the gene fusions of all unaligned reads from the PCAWG BAM files with the parameters of “MinimalHit=4, OutputFusionReads=True, RnaMode=True, FileFormat= BAM”. The FusionCatcher package was used to detect the gene fusions of raw paired-end RNA-seq reads, the option of “-U True; -V True” was set. Then only the fusions detected by two packages simultaneously were included in the final gene fusion set.

### Identification of SVs

The SVs were generated using both the SpeedSeq^85^ (release of v0.1.2) and GenomeSTRiP^86^ (release of v.2.00.1833) pipeline. SpeedSeq performed sample-level breakpoint detection via

LUMPY^87^ (release of v.0.3.1) followed by population-scale merging and genotyping of SV calls by svtools^88^ (release of v0.5.1). For GenomeSTRiP, we limited GSCNQUAL to one or more for detecting deletions and to eight or more for multiallelic CNVs, and to 17 or more for GenomeSTRiP duplications. The calling results of LUMPY and GenomeSTRiP were merged, and the duplicates were removed. Then we used MELT^89^ (release of v2.2.2) with the MELT-SPLIT function to identify Alu, SVA, and LINE-1 insertions.

### Identification of SNPs

The SNPs were detected by VarScan^90^ (release of v2.2.6) with parameters of

“--min-coverage 3 --min-var-freq 0.08 --p-value 0.10 --somatic-p-value 0.05 --strand-filter 0”, SomaticSniper^91^ (release of v1.0.4) with parameters of “-q 1 -Q 20”, and Strelka^92^ (release of v2.9.10) with the bwa_default parameters, and the extra parameter

“--ignore-conflicting-readnames”. All the detected SNPs were screened to remove common germline SNPs from a panel of whole-genome normal present at ≥ 0.1% MAF in dbSNP-138. All SNPs calls were required to satisfy maximum mismatch quality sum of ref-supporting reads ≤ 60; 5-bp maximum length of flanking homopolymer; minimum average relative distance from either end of reads ≥ 0.1; VAF < 2% with < 2 variant reads in the normal; minimum average relative distance to the effective 3’-end of read for var-supporting reads ≥ 0.2.

### The two-phase scoring algorithm: 3DFunc

To measure the effect of variants under the 3D context quantitatively, we proposed a two-phase scoring algorithm, 3DFunc, which combined the gene expression data from PCAWG and GTEx, as well as the Hi-C matrix from 4DN. To measure the gene expression changes between cancer and normal cell lines, the putative target gene *i* of predicted variants were mapped to the PCAWG expression data with the unique “aliquot”, the average matched gene expression of target gene *i* in *n*_*cancer*_ cancer cell lines was 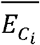, the standard deviation was 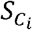, and the average matched gene expression of target gene *i* in *n*_*normal*_ normal cell lines was 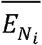, the standard deviation was 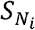. In this study, we used 20 cancer and 15 normal cell lines for calculation. The expression change between cancer and normal cell lines was calculated with t-test, which can be represented by,

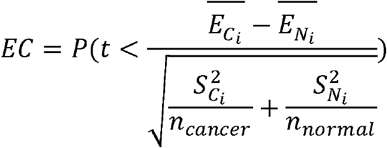

To characterize the 3D interaction frequency between predicted variant and target gene, we calculated the flexible IF for each variant-gene pair in 33 cell lines. For putative target gene *i*, the sum of flexible IF in 20 cancer cell lines was 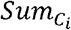, and the sum of flexible IF in 13 normal cell lines was 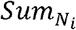, the flexible IF change was defined as 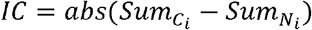.

Then we employed nonlinear least square curve^93^ to fit the expression change and flexible IF change of variant-gene pair. We observed that the data probably follow an exponential decay, thus we define the fitting function as *EC* = *Ae* ^−*IC*/τ^. In which the parameters of*A* and τ were determined by the residual sum of squares (RSS), since the goal of nonlinear least squares fitting algorithm is to find function parameters that minimize the RSS, which can be represented by *RSS* = ∑(*obs* − *pred*)^2^. The fitting function was implemented by homemade R script, the fitting result was stored as an R object named “model”. The 3DFunc score was then calculated by the difference between expected *EC* and observed *EC*.

### Scoring of Caviar Probability

To compare the scoring results of 3DFunc with other causal variants detecting methods, we calculated the Caviar Probability of our predicted variants with CAVIAR^57^, which took GWAS and eQTL linkage disequilibrium (LD) files as input to detect the causal variants, the Colocalization posterior probability (CLPP) for each variant was used as the final Caviar Probability.

### Construction of GAD Network

We firstly filtered the variant-gene pairs with the threshold of 3DFunc > 0.5 and p-value < 0.05, then extracted 100, 76, 116, 166, and 163 target genes from different blood, colon, liver, lung, and uterus individually. The GAD network was built with DisGeNET cytoscape app^94^ with the curated source, all the annotation types and all the disease class. The score and EI threshold were set from 0.5 to 1.0. The output associations included 831, 603, 971, 1091, and 925 diseases for each tissue. The GAD network was grouped by the type of variants, and the nodes were re-sized by the detected evidence. Finally, the diseases related to the accordingly tissues were selected and marked on the GAD network.

### Analysis of TCGA MDS data

For the TCGA MDS data, two cancer samples and two normal samples in BEATAML1.0-COHORT were used, the gene counts have been normalized by fragments per kilobase of exon per million mapped fragments (FPKM), which were taken as the input of DESeq2^95^ and used the default parameters to detect differentially expressed genes. Then p-value < 0.5 and the absolute value of log2(fold change) larger than or equal to 1 were used as the significant threshold to filter the genes, which showed 85 genes significantly differential expressed.

### 3DFunc of four common diseases

For heart tissue, we collected the RNA-seq and Hi-C data from samples at two critical time points during human embryonic stem cells (hESCs) differentiation: hESC (Day 0) and hESC-ventricular cardiomyocytes (Day 80) in the study of Zhang et al ^96^, under the GEO accession number of GSE116862. For pancreatic islet tissue, the RNA-seq data of four individuals were used ^97^ under the GEO accession number of GSE51312, and the Hi-C data were collected from the islets of 3 non-diabetic individuals under the AMP-CMD accession number of TSTSR043623. For breast tissue, the Hi-C and RNA-seq data were derived from MCF-10A mammary epithelial and MCF-7 breast cancer cell lines, under the GEO accession number of GSE66733 and GSE71862. For intestine tissue, the RNA-seq data were collected from IBD patient (12 samples) and healthy controls (6 samples) under the GEO accession number of GSE179128, the Hi-C data was available in the ArrayExpress database under accession E-MTAB-2323^98^. The RNA-seq and Hi-C data of each tissue were input to 3DFunc individually with the default extensive fold of 5, and chose the parameter of “pair_db” to get the calculated scores for the variant-gene pairs.

### Detection of gene list enrichment with DisGeNET

Gene list enrichments are identified in the ontology categories of DisGeNET ^58^. All genes in the genome have been used as the enrichment background. Terms with a p-value < 0.05, a minimum count of 3, and an enrichment factor > 1.5 (the enrichment factor is the ratio between the observed counts and the counts expected by chance) are collected and grouped into clusters based on their membership similarities. The top few enriched clusters (one term per cluster) are shown in the Figure 6. The algorithm used here is the same as that is used for pathway and process enrichment analysis in the Metascape web tool ^99^.

### Analysis of GPS according to 3DFunc percentile

Genome-wide polygenic scores (GPS) provide a quantitative metric of an individuals inherited risk based on the cumulative impact of many common polymorphisms. Weights are generally assigned to each genetic variant according to the strength of their association with disease risk. In the study of Amit et al. ^59^, the GPS weights of coronary artery disease (CAD), atrial fibrillation, type 2 diabetes, inflammatory bowel disease, and breast cancer were provided. Here we combined the GPS weights of CAD and atrial fibrillation as the cardiovascular disease for subsequent analysis. The variant-gene pairs were scored by 3DFunc in a disease-specific manner, then the pairs were grouped into 100 independent bins by calculating 1-100 percentiles thresholds of 3DFunc scores. And the average GPS weight was calculated for each bin to measure the clinical significance of 3DFunc scores.

### Fine-mapping scoring of EMS, PoPS, and ABC

The fine-mapping results of EMS in 49 GTEx tissues^60^ were downloaded from https://www.finucanelab.org/data, for each tissue, the list of top variant-gene pairs with highest EMS (normalized EMS > 100) were retained for subsequent analysis. And PoPS Score in 18 diseases with publicly available GWAS summary statistics and 95 complex traits from the UK Biobank^61^ were downloaded from https://www.finucanelab.org/data, the genomic coordinates of variants were transferred to hg38 with UCSC Liftover tool ^77^. The ABC scores in 72 diseases and complex traits^62^ were downloaded from https://www.engreitzlab.org/resources/, and the variant-gene pairs with ABC > 0.5 were retained. Then the variant-gene pairs from EMS, PoPS, and ABC were rescaled with z-score respectively, and the average rescaled value of EMS, PoPS, and ABC were calculated for each variant-gene pair as the final scaled score.

## Declarations

### Ethics approval and consent to participate

Not applicable

### Consent for publication

Not applicable

### Data availability

The GEO datasets used in this study are available under the accession numbers: GSE116862, GSE51312, GSE66733, GSE71862, and GSE179128. The ArrayExpress database under accession number of E-MTAB-2323, and AMP-CMD accession number of TSTSR043623. All the curated variant-gene pairs along with the 3D interruptions, as well as the 3DFunc scoring results are available at the 3DGeOD database: https://www.csuligroup.com/3DGeOD/home.

### Code availability

The source code of 3DFunc is available at GitHub repository: https://github.com/CSUBioGroup/3DFunc.

### Authors’ contributions

L.T. and M.L. conceived of the presented idea. L.T., M.H. and J.C. collected the data and designed the model, L.T. wrote the source code of 3DFunc. M.H. and J.C. developed the framework of database. M.C.H. helped improve the bioinformatics analysis. M.C.H., M.L. and P.T.E. aided in interpreting the results and provided input on the data presentation. All authors provided critical feedback and helped shape the research, analysis and manuscript.

## Acknowledgements

We are grateful to the High-Performance Computing Center of Central South University for partial support of this work.

